# Accounting for ambiguity in ancestral sequence reconstruction

**DOI:** 10.1101/409029

**Authors:** A. Oliva, S. Pulicani, V. Lefort, L. Bréhélin, S. Guindon

**Affiliations:** LIRMM, CNRS & Université de Montpellier, Montpellier, France; Australian Centre for Ancient DNA, Adelaide, Australia

## Abstract

The reconstruction of ancestral genetic sequences from the analysis of contemporaneous data is a powerful tool to improve our understanding of molecular evolution. Various statistical criteria defined in a phylogenetic framework can be used to infer nucleotide, aminoa-cid or codon states at internal nodes of the tree, for every position along the sequence. These criteria generally select the state that maximises (or minimises) a given criterion. Although it is perfectly sensible from a statistical perspective, that strategy fails to convey useful information about the level of uncertainty associated to the inference. The present study introduces a new criterion for ancestral nucleotide reconstruction that selects a single state whenever the signal conveyed by the data is strong, and a combination of multiple states otherwise. Simulations demonstrate the benefit of this approach with a substantial increase in the accuracy of ancestral sequence reconstruction without significantly compromising on the precision of the solutions returned.

## Introduction

Molecular sequences collected in present day species provide a wealth of information about past evolutionary events. Using relevant probabilistic models of molecular evolution, it is possible to reconstruct the sequences of species ancestral to a sample of taxa. The application of these techniques led to spectacular findings. In particular, the resurrection of ancestral proteins using biochemical processes (*Chang and Donoghue, 2000, Thornton, 2004, Bridgham et al., 2006*) helped improved our understanding of the ways evolution takes place at the molecular level.

Phylogenetics provides an adequate framework for the reconstruction of ancestral sequences. Given a phylogenetic tree that depicts the evolutionary history of a sample of taxa along with a set of corresponding homologous sequences, it is possible to estimate the sequences at each internal node of the tree. The parsimony approach (*Fitch, 1971*) consists in selecting the ancestral sequences that minimise the number of substitutions required to explain the sequences observed at the tips of the tree. It uses information related to the grouping of taxa in the tree while amounts of evolution (i.e., the length of edges in the tree) are ignored. The accuracy of the ancestral sequences estimated with the parsimony approach has been well studied from a theoretical perspective (*Fischer and Thatte, 2009, Yang et al., 2011*) and using simulations (*Zhang and Nei, 1997*).

(*Yang et al., 1995*) and (*Koshi and Goldstein, 1996*) were the first to infer ancestral sequences using the maximum likelihood approach under a phylogenetic model. The inference relies here on a twostep approach. A phylogenetic model (i.e., a tree topology, with edge lengths and substitution model parameters) is first fitted to the data. Next, finding the nucleotide (or amino-acid or codon) character with the highest (marginal) posterior probability at any internal node in a fixed phylogeny is relatively straightforward. (*Pupko et al., 2000*) later described an elegant dynamic programming algorithm for the inference of the combination of character states at all internal nodes that maximises the joint posterior probability.

Yet, these methods focus on selecting the best state in the alphabet of actual nucleotides, amino-acids or codons, i.e., the alphabet the data generating process relies upon. Although it is perfectly sensible from a statistical viewpoint, it does not always reflect potential uncertainty in the inference. Indeed, although multiple characters can have high probabilities, selecting only the best one obliterates available information about the variability of ancestral sequence estimates. Ignoring uncertainty in the reconstruction of ancestral sequences is indeed a serious limitation of current estimation techniques that has been known for a long time (*Cunningham et al., 1998*).

In the present study, we introduce a new statistical criterion, the minimum posterior expected error (MPEE), that reveals uncertainty in the inference without compromising on the interpretability of ancestral character reconstruction. We focus on the inference of past DNA sequences. In this context, we show that accommodating for ambiguity in ancestral nucleotides using this new criterion amounts to a better use of the available data compared to the traditional approaches.

### Notations

The multiple sequence alignment is noted as **d**. Its length, i.e., the number of columns in the alignment is *l*. Each sequence in **d** is a string of characters, each character being taken within the alphabet 𝒜 of nucleotides, amino-acids, codons or any other well-defined discrete states in finite number. In what follows, **d**^(*s*)^ corresponds to the *s*-th column of **d**. Let *τ* denote the unrooted topology of the phylogeny under scrutiny and **l** the vector of edge lengths on that tree. We assume that the tree is binary so that there are *n* – 2 vertices, where *n* is the number of tips. Let x denote the vector of internal nodes and *x* is one of these nodes. *v*_1_(*x*), *v*_2_(*x*) and *v*_3_(*x*) are the three nodes directly connected to *x*, i.e., its three direct neighbours. 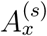 is the random variable giving the ancestral character observed at node *x* and site *s*.

Inferring ancestral characters generally relies on evaluating the conditional probability 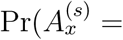 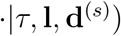, which, for a particular character state *y*, is expressed as follows:

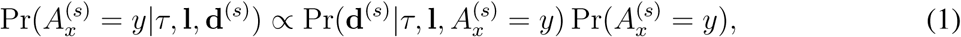

where 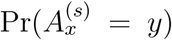 is taken as the equilibrium frequency of state *y* since the substitution process at each site of the alignment is modelled as a homogeneous Markov chain running along the phylogeny. 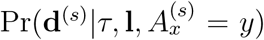 is the likelihood of the model given that state *y* is observed at node *x*. Assuming that node *x* has three neighbours, as is the case if the phylogeny is a fully resolved unrooted tree, 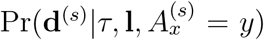 is obtained as follows:

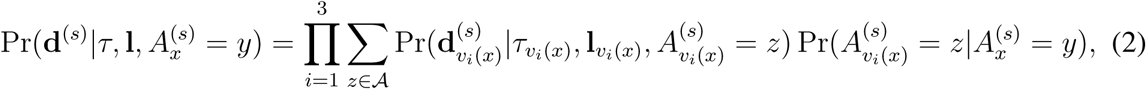

where **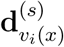** corresponds to the part of **d** made of sequences found at the tips of the subtree rooted by *v*_*i*_(*x*) (hence **d**_*v*_1_(*x*)_ ∪ **d**_*v*_2_(*x*)_ ∪ **d**_*v*_3_(*x*)_ = **d**). *τ*_*v_i_*_(*x*) is the topology of this rooted subtree and **l**_*v_i_*(*x*)_ are the lengths of its edges.

### Inferring ancestral states

In the following two sections, we introduce the standard criterion, i.e., the maximum a posteriori criterion, defined in the context of an unrooted or a rooted tree. The new, minimum posterior expected error, is presented next.

### The maximum a posteriori (MAP) criterion

The most popular technique to infer ancestral states relies on the maximum a posteriori probability (MAP) criterion based on the marginal conditional probabilities defined above. The inferred character *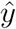* is selected as follows:

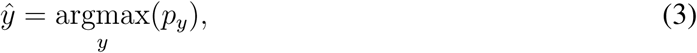

Where 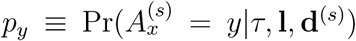. This technique thus considers each node separately,selecting the most probable state at each of these nodes without any reference to the ancestral states inferred in other parts of the tree. As mentionned earlier, dynamic programming can also be used to infer the combination of ancestral states at all internal nodes that maximizes their joint posterior probability (*Pupko et al., 2000*). In practice however, most if not all phylogenetic software rely on the marginal probabilities.

In Equation 3, the marginal posterior probability derives from the conditional likelihood as defined in Equation 2. Since the sum in this last equation is over the three vertices directly connected to the internal node under scrutiny, the whole set of characters found at the tips of the tree is involved in the calculation of this probability. A distinct approach, which applies to the case where the tree is rooted, is to sum over the two vertices subtending the node under scrutiny only (i.e., the two nodes “away from the root”, taking the node of interest as reference). This solution amounts to considering that only the data “below” the node of interest convey information about the ancestral state. This reasoning is somehow at odds with the time-reversible Markovian assumption about the substitution process and unnecessarily ignores part of the data. This approach is nonetheless implemented in the software RAxML (*Stamatakis, 2006*). In the following, we will refer to this approach as the MAP_r_ criterion.

### The minimum posterior expected error (MPEE) criterion

The MPEE criterion is closely related to the MAP approach in particular settings. The rationale behind MPEE rests on the definition of the loss function that the estimation of an ancestral character relies upon. The simplest way to define such function is as follows:

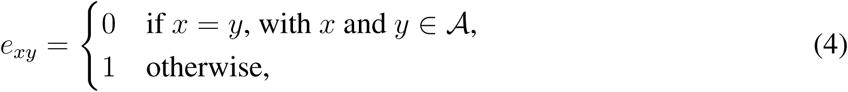

where *e*_*xy*_ is the error score when comparing *x*, the true character and *y* the inferred one. It is equal to 0 when the inferred and the true states are the same and 1 otherwise.

The posterior expected error for an inferred character *y* is a weighted average of these errors. It is obtained as follows:

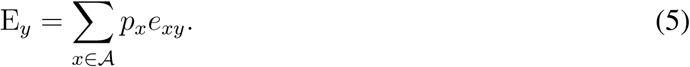

when the loss function is as defined in Equation 4, the posterior expected error is E_*y*_ = 1 – *p*_*y*_. Hence, in this particular setting, selecting the character that minimizes the posterior expected error, i.e., finding *y*^*\**^ such that *y*^*\**^ = arg min_*y*∈𝒜_ (E_*y*_), is the same as selecting the character that maximises the posterior probability.

In general however, it is of interest to consider an alphabet of inferred states, **𝒜***’*, that is distinct from that of the actual states involved in the data generating process, **𝒜** (see Table 1). The posterior expected error can still be evaluated using Equation 5. However, the criterion that is used for the inference is now *y*^*\**^ = arg min_*y*∈𝒜’_ (E_*y*_). Since there is no guarantee that arg min_*y*∈𝒜_ (E_*y*_) = arg min_*y*∈𝒜’_ (E_*y*_) in general (but see below), the character selected using the MPEE criterion might differ from that selected using MAP.

**Table 1:** Cost of errors associated to each non-ambiguous and ambiguous state. The rows give the list of characters the MPEE criterion can choose from. The columns give the actual nucleotide state the data generating process relies upon. Each element of the table gives the cost associated to the corresponding pair of (inferred and true) character states.

| $x$ | $\{A\}$ | $\{C\}$ | $\{G\}$ | $\{T\}$ |
| --- | --- | --- | --- | --- |
| $y$ | | | | |
| $\{A\}$ | $\alpha_1$ | $\beta_1$ | $\beta_1$ | $\beta_1$ |
| $\{C\}$ | $\beta_1$ | $\alpha_1$ | $\beta_1$ | $\beta_1$ |
| $\{G\}$ | $\beta_1$ | $\beta_1$ | $\alpha_1$ | $\beta_1$ |
| $\{T\}$ | $\beta_1$ | $\beta_1$ | $\beta_1$ | $\alpha_1$ |
| $R:=\{G,A\}$ | $\alpha_2$ | $\beta_2$ | $\alpha_2$ | $\beta_2$ |
| $Y:=\{C,T\}$ | $\beta_2$ | $\alpha_2$ | $\beta_2$ | $\alpha_2$ |
| $S:=\{G,C\}$ | $\beta_2$ | $\alpha_2$ | $\alpha_2$ | $\beta_2$ |
| $W:=\{A,T\}$ | $\alpha_2$ | $\beta_2$ | $\beta_2$ | $\alpha_2$ |
| $K:=\{G,T\}$ | $\beta_2$ | $\beta_2$ | $\alpha_2$ | $\alpha_2$ |
| $M:=\{A,C\}$ | $\alpha_2$ | $\alpha_2$ | $\beta_2$ | $\beta_2$ |
| $B:=\{C,G,T\}$ | $\beta_3$ | $\alpha_3$ | $\alpha_3$ | $\alpha_3$ |
| $D:=\{A,G,T\}$ | $\alpha_3$ | $\beta_3$ | $\alpha_3$ | $\alpha_3$ |
| $H:=\{A,C,T\}$ | $\alpha_3$ | $\alpha_3$ | $\beta_3$ | $\alpha_3$ |
| $V:=\{A,C,G\}$ | $\alpha_3$ | $\alpha_3$ | $\alpha_3$ | $\beta_3$ |
| $N:=\{A,C,G,T\}$ | $\alpha_4$ | $\alpha_4$ | $\alpha_4$ | $\alpha_4$ |

Table 1 defines the loss function when the inferred nucleotide characters are not systematically non-ambiguous. The definition is incomplete here since the values of the parameters *α*_1_, *…, α*_4_ and *β*_1_, *…, β*_3_ are not set yet. In order to do so, we rely on the definition of the mean prior error for a given character *y* ∈ 𝒜’ as follows:

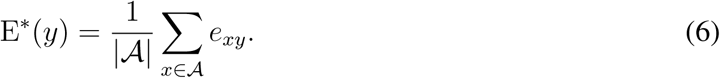

E^*\**^(*y*) is thus simply a weighted average error with the weight of each character in 𝒜 equal to 1/|𝒜|. Because there is no obvious reason that a character in 𝒜’ should have a mean prior error that differs from that obtained for another character, we choose to have E^*\**^(*y*) = E^*\**^(*y’*) for any *y* and *y*’ ∈ 𝒜’. This last equality helps us define a simple linear relationship between *α*_1_ and *β*_1_. Indeed, for any *y* ∈ 𝒜, we have:

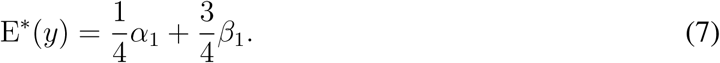

In the present study, we choose to have E^*\**^(*y*) = 3/4 which is the value of E^*\**^(*y*) obtained in the special case where *α*_1_ = 0 and *β*_1_ = 1.

We now determine the expressions of *α*_2_ and *β*_2_ such that the mean prior error for any *y* ∈ {R,Y,S,W,K,M}, i.e., for doubly-ambiguous states, is also equal to 3/4. In order to do so, we need to set the values of *α*_2_ and *β*_2_ such that the following equation holds:

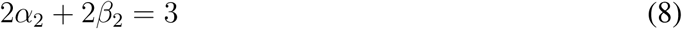

An infinite number of values of *α*_2_ and *β*_2_ satisfy this equality. We define a range of relevant values for *α*_2_ (and *β*_2_) as follows. A first rationale is to have *α*_2_ = 0 (and thus *β*_2_ = 3/2) so that the error when inferring an ambiguous character that is compatible with the true state (e.g. *y* = ‘R’ while *x* = ‘A’) is set to zero, in a similar manner to what is done when the inferred state is not ambiguous. Another reasoning would be to have *β*_2_ = 1 (and thus *α*_2_ = 1/2) so that the error when inferring an ambiguous character that is not compatible with the true state (e.g., *y* = ‘Y’ while *x* = ‘A’) is equal to one. Since there is no obvious reason to select one option or the other or any particular choice of *α*_2_ (or *β*_2_) in between, we considered a range of multiple values of *α*_2_ in the 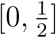 interval in our simulations.

Having a mean prior error equal to 3/4 for any triply-ambiguous state means that the following relationship also holds:

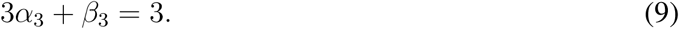

Using the same rationale as before, having *α*_3_ = 0 (and *β*_3_ = 3) and *β*_3_ = 1 (and *α*_3_ = 2/3) define a range of relevant values for these two parameters. In our simulations, we thus considered multiple values of *α*_3_ in 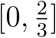.

Let *p*_(1)_, *…, p*_(4)_ be the ordered marginal probabilities of states in 𝒜 such that *p*_(1)_ *≥ p*_(2)_ *≥ p*_(3)_ *≥ p*_(4)_. In the special case where *α*_1_ = *α*_2_ = *α*_3_ = 0, the posterior mean errors that are minimal among the non-ambiguous as well as the doubly- and triply-ambiguous states are respectively equal to:

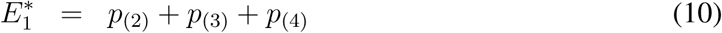

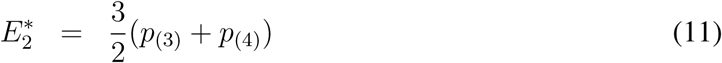

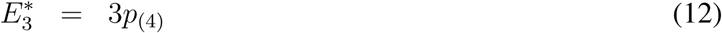

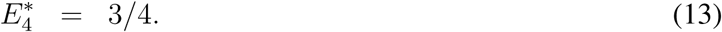

The mean posterior error will thus systematically be minimized by a triply-ambiguous case (except in the special situation where *p*_(1)_ = *p*_(2)_ = *p*_(3)_ = *p*_(4)_ = 1/4 in which case all posterior mean errors are equal to 3/4). Also, when *β*_1_ = *β*_2_ = *β*_3_ = 1, we have

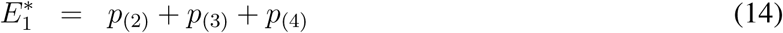

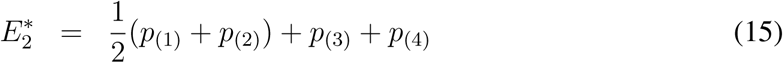

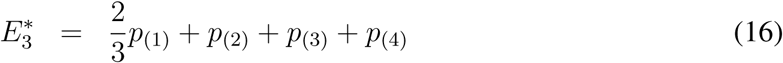

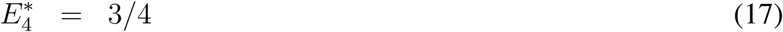

In this special case, we have *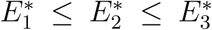* and *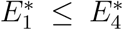* such that the posterior mean error will systematically be minimised by a non-ambiguous state. Hence, the cases where *α*_2_ = *α*_3_ = 0 and *β*_2_ = *β*_3_ = 1 correspond to two extremes where the level of ambiguity of the state minimising the criterion is fully determined by the values of these parameters.

As already stated above, it is not obvious which values of *α*_2_ and *α*_3_ should be chosen between these two extremes. We thus considered a grid of multiple values for these two parameters. The inferred characters was then taken as the most frequent among all states inferred using particular values of *α*_2_ and *α*_3_. More specifically, we used a grid of 50 equally spaced values of *α*_2_ in [0, 1/2] by 50 equally spaced values in [0, 2/3] for *α*_3_ with the constraint that *α*_3_ *≥ α*_2_. This last inequality guarantees that the cost of a match obtained with a triply-ambiguous states is greater or equal to that obtained with a doubly-ambiguous character. Results (not shown) obtained with a higher mesh density gave results very similar to that derived with 50 *×* 50/2 points only.

For particular values of *α*_2_ and *α*_3_, a naive approach to identify the character (ambiguous or not) that minimises the posterior mean error would be to evaluate this error for all states. Yet, the computational complexity can be considerably reduced by examining closely the error in each class of ambiguity. For non-ambiguous states, the minimum of the posterior mean error is indeed equal to

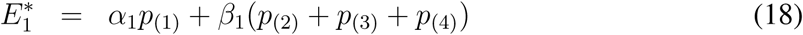

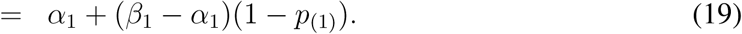

Hence, when considering non-ambiguous characters only, the state that minimises the posterior mean error is, unsurprisingly, the one with the highest posterior probability. Now, for doubly-ambiguous state, the minimum is equal to

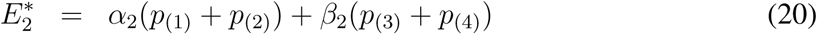

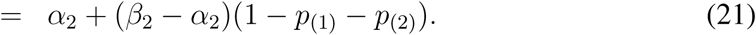

Therefore, the optimal pair of non-ambiguous states is the one with the highest and second highest posterior probabilities associated to the two (non-ambiguous) states making up the pair. Similarly, for triply-ambiguous states, we have:

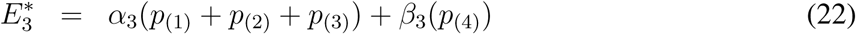

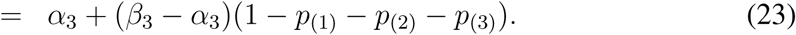

Here again, finding the best triplet simply requires identifying the three most probable non-ambiguous states. Hence, for particular values of *α*_2_ and *α*_3_, the time complexity for computing *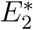* and 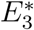 is 𝒪(|𝒜|) instead of 𝒪(|𝒜|^2^) and 𝒪(|𝒜|^3^) respectively when calculations are conducted in a naive fashion. Our approach is therefore not limited to the analysis of nucleotide data and amino-acid or even codon data can be accommodated too.

The MPEE criterion is not the only one that can be used to achieve the same goal of inferring ambiguous or non-ambiguous ancestral states. Very recently, Ishikawa and colleagues (*Ishikawa et al., 2018*) described a method for reconstructing ancestral sequences based on the Brier scoring rule. This approach consists in comparing the observed distribution of posterior probabilities obtained for non-ambiguous states to that of null (or expected) uniform distributions on 1, 2, …, |𝒜| states. A Euclidean distance characterises the difference between the observed and each of the |𝒜| expected distribution. The inferred state, which can be ambiguous or not, is the one that minimises these Euclidean distances. Both approaches are similar in spirit and have the same time complexity. Yet, as opposed to Ishikawa’s approach, the MPEE criterion does not rely on specifying null distributions of posterior probabilities for the different levels of ambiguity. Hence, in theory at least, there is no obvious reason that suggests that Ishikawa’s method could outperform MPEE.

### Simulations

The performance of several software and criteria used in the estimation of ancestral sequences was assessed using multiple simulated data sets. Each such data set consists in a phylogenetic tree along which nucleotide sequences are evolved according to a standard Markov model of sequence evolution. The specifics of these simulations are given below.

### Simulating phylogenies

Random trees were generated with the R package *TreeSim* (*Stadler, 2010*). *TreeSim* creates random trees under a constant rate birthdeath process. The function *sim.bd.taxa.age* was used to generate trees with a fixed number of taxa and fixed time since the most recent common ancestor of the sampled species. The number of taxa in the generated tree is equal to 50. The tree height parameter (parameter ‘`age`’ in `sim.bd.taxa.age`), *H*, was randomly drawn for each tree in *U* [0.1; 1]. 50 trees were generated in total. The birth rate was fixed to 0.1 while the extinction rate was fixed at 0.5. The trees hence generated are utltrametric, i.e., clocklike. The edge lengths are interpreted here as amounts of nucleotide substitutions per site with an average length equal to 0.08. We kept the rooted version of the trees along with the corresponding unrooted trees. The code that was used to generate and prepare the trees is accessible from https://gite.lirmm.fr/atgc/phyml/AncestralSequences_Oliva2018/blob/master/Create_Tree.R.

### Simulating sequences

We used the software *INDELible* (*Fletcher and Yang, 2009*) to simulate nucleotide sequences along the trees. *INDELible* evolves sequences under a probabilistic model of molecular evolution that accommodates for point substitutions. It also accounts for insertion and deletion events although we did not consider this option in the present study. The GTR model (*Tavaré, 1986*) was used with rates of substitutions drawn in a uniform distribution in the [0.5, 1.5] interval for transitions and in a normal distribution with mean set to 4.0 and standard deviation equal to 4.0 for transversion. This normal density was truncated so that only positive rate values are considered. Nucleotide frequencies were drawn randomly in a uniform distribution in [0, 1] and normalised. Also, the length of the ancestral sequence at the root of the tree was set to 1,000 bp for all simulations. The scripts that were used to generate sequences are available at the following address: https://gite.lirmm.fr/atgc/phyml/AncestralSequences_Oliva2018/blob/master/Scripts.py.

### Computer programs and parameter settings

For each simulated data set, we inferred ancestral sequences using the same nucleotide substitution model as that used for the simulations (i.e., GTR) and a simplified model (*Jukes and Cantor, 1969*), noted as JC in the following. The parameters of the GTR model were estimated using maximum likelihood under the “true”, i.e., the simulated phylogeny. Also, ancestral sequences were inferred using this tree so that it is straightforward to match ancestral sequences to the corresponding estimated ones.

PhyML (*Guindon et al., 2010*) and RAxML (*Stamatakis, 2006*) were used to reconstruct ancestral sequences under the JC and GTR models. While the tree topologies used for the estimation were set to the true ones and fixed throughout the analysis, edge lengths and substitution model parameters (for the GTR model) were optimised with each software prior the ancestral sequence reconstruction.

## Results

Figure 1 gives the fraction of errors in ancestral state inference when using the MAP, MAP_r_ and MPEE criteria as a function of the difficulty of the estimation. The difficulty is quantified here based on the value of the maximum marginal probability, *p*_(1)_. For instance, inference is more prone to errors in cases where *p*_(1)_ is close to 0.25 compared to cases where it is close to 1.0. We therefore considered a range of threshold values where the fraction of errors was measured only in cases where the maximum marginal probability was below this threshold. The fraction of errors is straightforward to measure in cases where the inference returns only non-ambiguous states. Since inferred states can be ambiguous when using MPEE, we used different approaches to measure the errors here. We first considered that any state, ambiguous or not, compatible with the true one is associated to an error cost equal to zero, while any non-compatible state has cost one (plain blue line in Figure 1). We also considered the case where *α*_2_ = *α*_3_ = 0, *β*_2_ = 3/2 and *β*_3_ = 3, i.e., the cost of compatibility (i.e., zero) is the same for non-ambiguous and ambiguous states, while the cost of incompatibility is equal to 3/2 for doubly-ambiguous states and 3 for triply-ambiguous states (dashed blue line). Finally, we considered the case where *β*_2_ = *β*_3_ = 1, *α*_2_ = 1/2 and *α*_3_ = 2/3, i.e., the cost of incompatibility (i.e., one) is the same for ambiguous and non-ambiguous inferred states while the cost of compatibility is zero for non-ambiguous states, 1/2 for doubly-ambiguous and 2/3 for triply-ambiguous states (dotted blue line).

**Figure 1:**
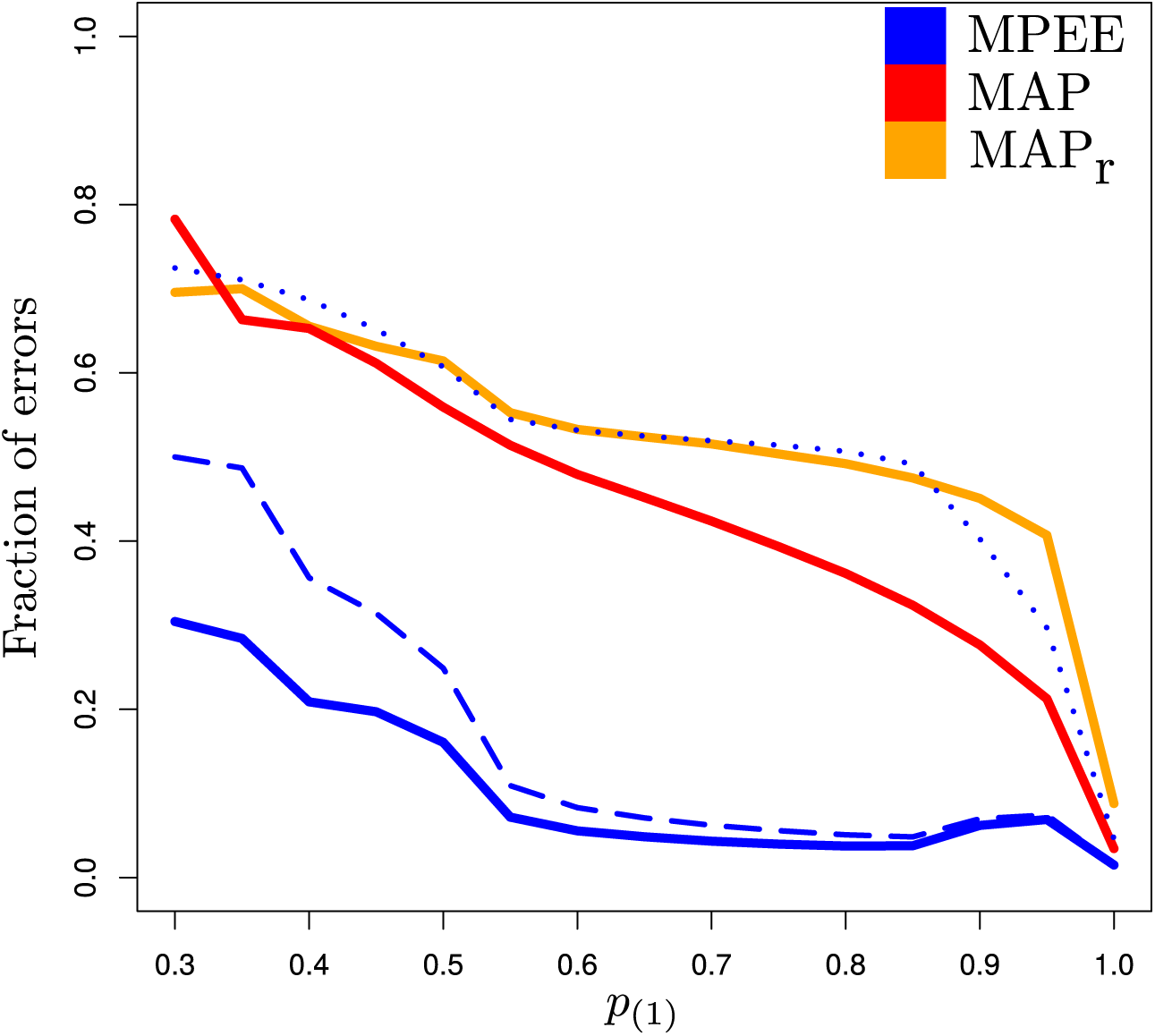
Accuracy of ancestral state reconstruction when using the MAP, MAP_r_ and MPEE criteria (the GTR model was used for the inference). The horizontal axis gives the values of the maximum marginal probabilities below which accuracy is measured (see text). The vertical axis gives the fraction of errors, i.e., the proportion of cases where the inferred ancestral state is not compatible with the actual (i.e., true) state. The fraction of errors is measured in three different ways for MPEE: (1) an incompatible state incurs a cost of one while a compatible one costs zero (plain line); (2) an incompatible state is associated to a cost of one while a compatible one costs 0, 1/2 or 1/3 depending on whether the state is non-ambiguous, doubly-ambiguous or triply-ambiguous respectively (dotted line); (3) an incompatible state costs 1, 1.5 or 3 whether it is non-ambiguous, doubly-ambiguous or triply-ambiguous respectively while a compatible state costs 0 (dashed line).

Results in Figure 1 first show that the MAP_r_ underperforms compared to other criteria. Hence, considering the characters states that lie at the tips below a given node in a rooted tree and ignoring all the nucleotides at the tips elsewhere is, perhaps unsurprisingly, prone to more errors than when using all the data available for the inference. The fraction of errors obtained with MAP and MPEE are generally smaller than that of MAP_r_.

The most accurate results are obtained with MPEE when the ambiguous status of the inferred states does not influence the way errors are measured (i.e., any incompatible state has an error cost of one while the cost is equal to zero otherwise). When the cost of compatibility depends on the level of ambiguity (0 for non-ambiguous, 1/2 for doubly-ambiguous and 2/3 for triply-ambiguous), then the fraction of errors is close to that obtained with MAP_r_ (see dotted line). If instead, it is the cost of incompatibility that depends on the level of ambiguity (1 for non-ambiguous, 3/2 for doubly-ambiguous and 3 for triply-ambiguous), then the fraction of errors is low (see dashed blue line). Hence, the accuracy with which MPEE reconstructs ancestral sequences very much depends on how one measures the error. The interpretation of results will indeed vary substantially whether one elects to heavily penalise errors when ambiguous states are inferred (*β*_2_ = 3/2 and *β*_3_ = 3) or whether one chooses to penalise correct inferences because of the ambiguous status of the inferred states (*α*_2_ = 1/2 and *α*_3_ = 2/3).

Figure 2 displays the results obtained when using the JC substitution model for the inference instead of the (correct) GTR model. The fraction of errors obtained here are similar to that observed when the substitution model is correctly specified. The mean error fraction when using the MAP_r_ and the MAP criteria are slightly higher under the JC model compared to GTR (0.44 *vs.* 0.43 for MAP_r_ and 0.40 *vs.* 0.36 for MAP) while MPEE is not affected at all (the fraction of errors is equal to 0.04 (plain blue line) in both cases). One also notes a substantial increase of the error fraction when the highest marginal probability is below 0.4, i.e., when the inference problem is particularly difficult. Hence, model misspecification impacts the quality of the inference mostly in cases where the signal conveyed by the data about ancestral sequences is weak.

**Figure 2:**
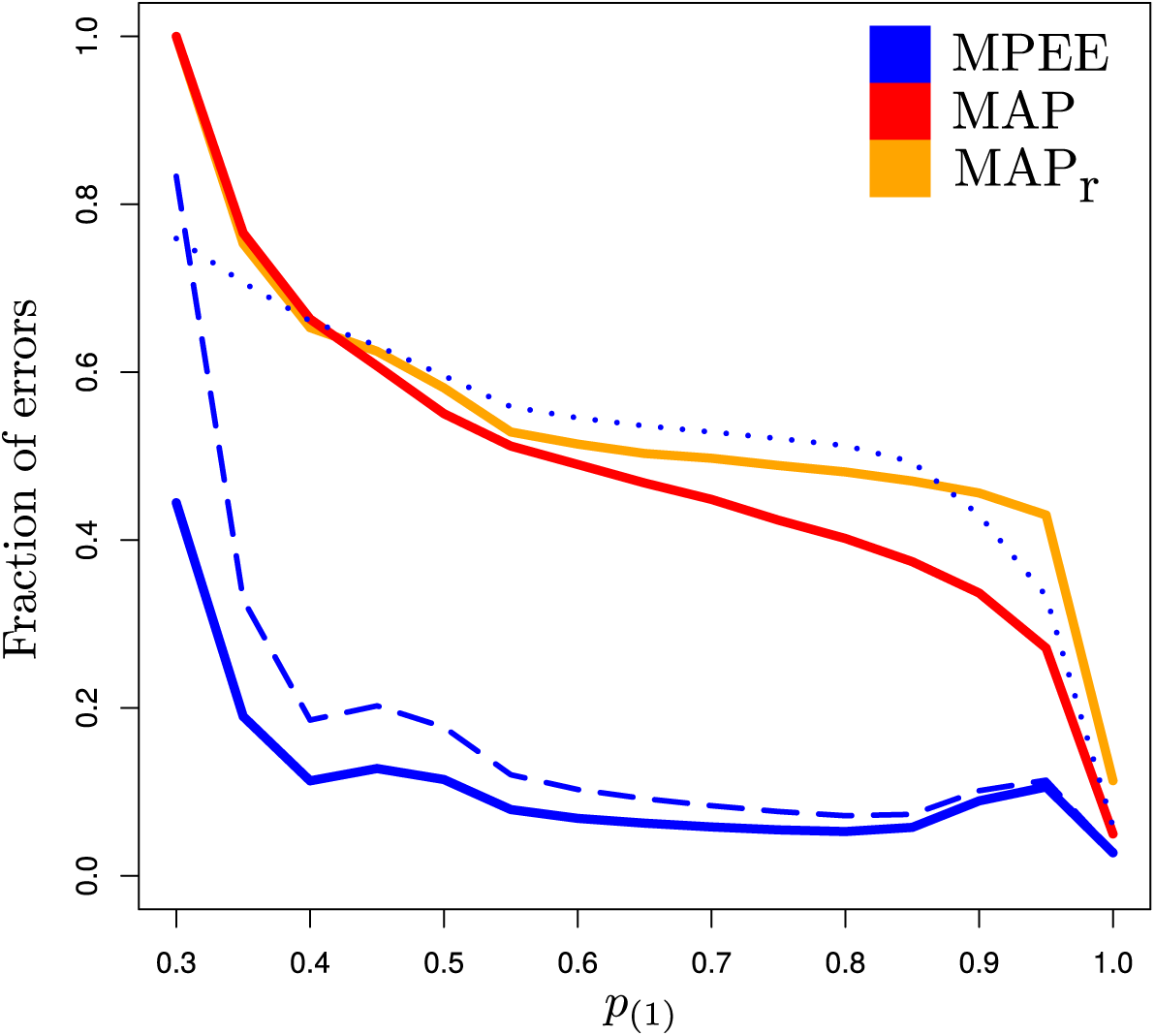
Accuracy of ancestral state reconstruction when using the MAP, MAP_r_ and MPEE criteria (the JC model was used for the inference). See caption of Figure 1.

We then focused on the distribution of the marginal probability of the states inferred by the MAP and the MPEE criteria. Examination of these distributions shows that the 1% least-probable inferred states have marginal probabilities equal to 0.8 and 0.6 with MPEE and MAP respectively when the inference is conducted under the GTR model of substitution. When the JC model is used instead the corresponding marginal probabilities are equal to 0.9 and 0.6 respectively. This result suggests that model misspecification does not affect the distribution of the ancestral state probabilities. Figures 3 and 4 display the quantiles of the two distributions (one for each criterion) corresponding to cumulative probabilities in the [0.0,0.2] interval for the GTR and the JC models. The similarity between the results obtained here with the two substitution models further confirms the robustness of the marginal probability estimates. The sharp increase of the marginal probabilities as obtained with MPEE in contrast to that observed with MAP indicates that virtually all the states selected by MPEE, including the least probable ones, reach a marginal probability close to 1.0 while that selected by MAP are less probable overall.

**Figure 3:**
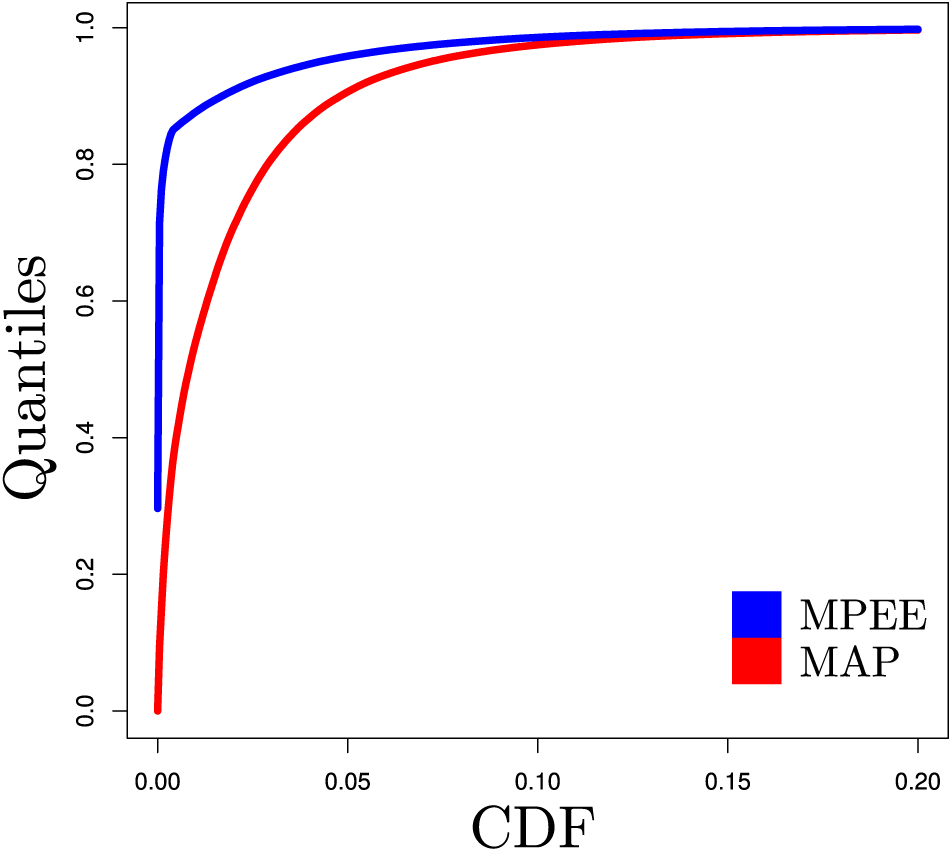
Quantiles of the marginal probability of states inferred using the MAP (in red) and MPEE (in blue) criteria, with the GTR model used for the inference. The vertical axis gives the values of *x* in Pr(*X ≤ x*), where *X* is the marginal probability of the inferred state. The horizontal axis gives the values of Pr(*X ≤ x*).

**Figure 4:**
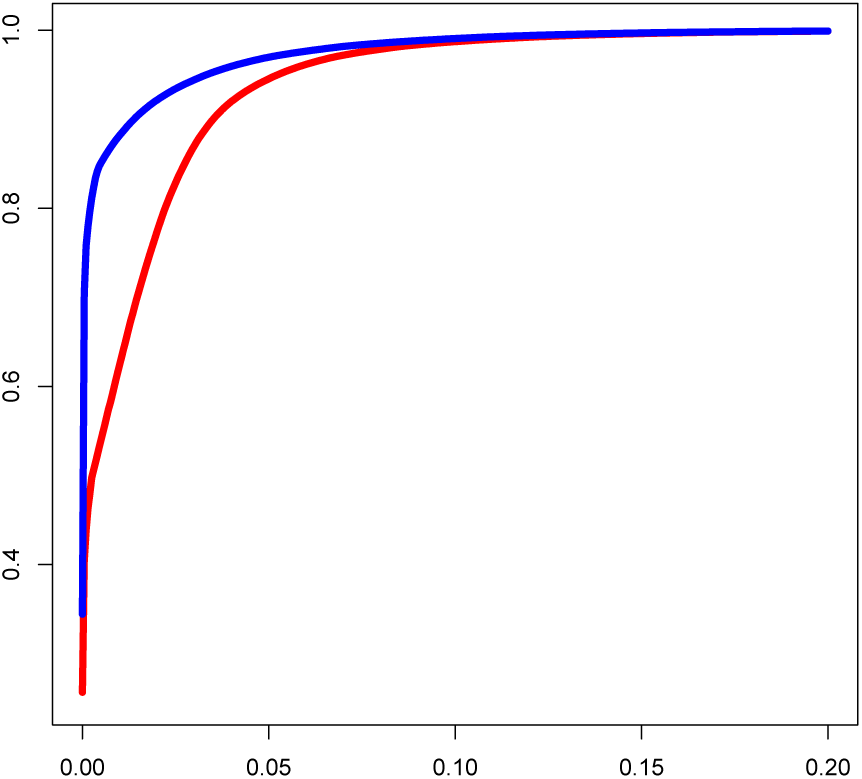
Quantiles of the marginal probability of states inferred using the MAP (in red) and MPEE (in blue) criteria, with the JC model used for the inference. See caption of Figure 3.

Table 2 gives the number of correct and incorrect inferred states selected using the MAP and the MPEE criteria, along with the ambiguity status of the character selected by MPEE when the model is correctly specified. Analysis of the figures presented in this table shows that the percentage of states incorrectly estimated using the MAP criterion that are correct when using MPEE is equal to 56%. In other words, more than half of the errors arising when using MAP can be corrected by applying the MPEE criterion. The MPEE criterion thus provides a substantial improvement in terms of the accuracy of the ancestral state reconstruction. Also, the fraction of ambiguous and correct states according to MPEE among the states correctly estimated using MAP, which we will refer to as precision, is equal to 2%. Therefore, the observed gain in accuracy entails little loss in terms of the precision with which ancestral states are reconstructed.

**Table 2:**
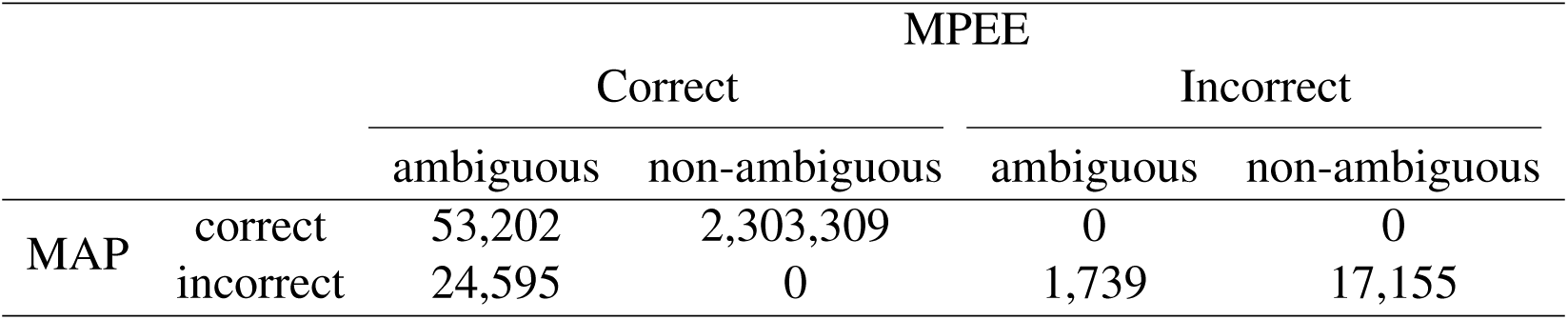
Number of correctly and incorrectly inferred ancestral states using the MAP *vs.* the MPEE criterion under the GTR model of substitution. Characters inferred using the MPEE criterion are also classified into ambiguous and non-ambiguous states.

Table 3 presents the results obtained when the substitution model is misspecified. The fraction of errors when applying the MAP criterion that are corrected using MPEE is now equal to 45%. Also, only 1.5% of the states correctly estimated using MAP are ambiguous (and correct) according to MPEE. Hence, model misspecification entails a decrease of the ability of MPEE to correct for errors arising when using MAP. Yet, this decrease of accuracy does not come with a decrease in terms of the precision of the estimates.

**Table 3:** Number of correctly and incorrectly inferred ancestral states using the MAP *vs.* the MPEE criterion under the JC model of substitution. See caption of Table 3.

|  |  | MPEE |  |  |  |
| --- | --- | --- | --- | --- | --- |
|  |  | Correct |  | Incorrect |  |
|  |  | ambiguous | non-ambiguous | ambiguous | non-ambiguous |
| MAP | correct | 35,907 | 2,317,662 | 0 | 0 |
|  | incorrect | 20,839 | 0 | 1,956 | 23,636 |

Finally, Figure 5 depicts the relationships between the MPEE and two other statistical criteria that may help better understand the circumstances in which MPEE selects a particular level of ambiguity. The first alternative criterion is *p*_(1)_, i.e., the maximum posterior probability of a non-ambiguous state. The rationale here is that the smaller *p*_(1)_, the more likely it is that MPEE will select an ambiguous state. The second criterion is the entropy of the posterior probabilities which is maximum whenever all probabilities are equal and minimum when a posterior probability is equal to one. The plot obtained shows that the relationship between the different criteria is not trivial. It is fairly clear that triply-ambiguous states are generally chosen whenever *p*_(1)_ is close to 1/𝒜 = 0.25 and/or the entropy is high. Yet, the level of ambiguity of the state selected by MPEE is not fully determined by threshold values applied to *p*_(1)_ or the entropy.

It is perhaps puzzling to observe instances where very similar values of the entropy and *p*_(1)_ lead to MPEE either choosing a doublyor a triply-ambiguous character. For instance, in case where *p*_(1)_ = 0.40, *p*_(2)_ = 0.33, *p*_(3)_ = 0.22 and *p*_(4)_ = 0.05, MPEE will select a doubly-ambiguous character. When we have *p*_(1)_ = 0.41, *p*_(2)_ = 0.30, *p*_(3)_ = 0.25 and *p*_(4)_ = 0.05, MPEE will choose a triply-ambiguous character instead. Although the values of *p*_(1)_ (0.40 and 0.41) and the entropy (1.75 and 1.76) are virtually identical in both cases, the difference between *p*_(2)_ and *p*_(3)_ is more important in the first case compared to the second one, thereby explaining the choices made by MPEE. Altogether, these results show that inferring the level of ambiguity based on thresholds of the entropy or *p*_(1)_ would lead to results different from that obtained with MPEE. We argue that these alternative criteria are less able than MPEE to extract relevant information from the data (i.e., the posterior probabilities).

## Discussion

This study introduces a new statistical criterion for the inference of ancestral nucleotide sequences. The main motivation behind our work was to improve the way uncertainty in the ancestral reconstruction is dealt with. In particular, in cases where multiple nucleotides have similar marginal probabilities, selecting only one of them can be problematic. Doing so is prone to an increased amount of errors in the inference (see lefthand sides of Figures 1 and 2). Also, this approach fails to convey important information about uncertainty in the inference.

**Figure 5:**
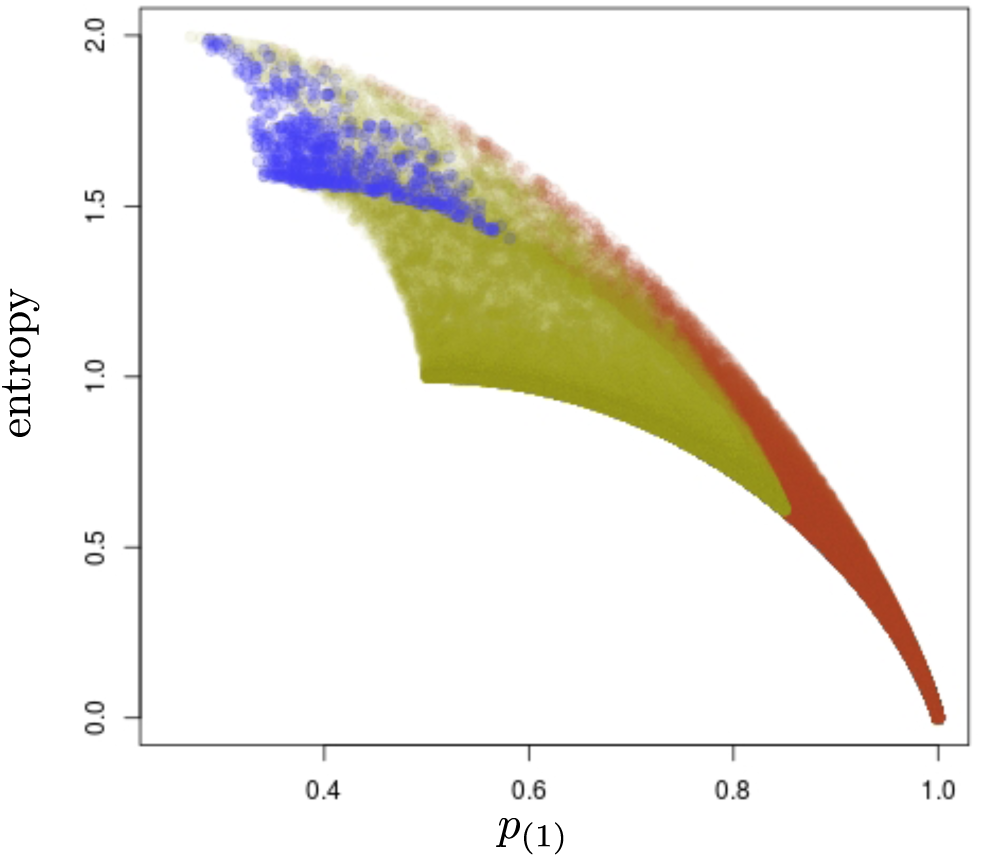
Relationship between MPEE, *p*_(1)_, and entropy. *p*_(1)_ is the maximum posterior probability evaluated at a given site and node in our simulations. The entropy is here obtained as *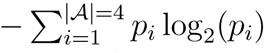*. It is equal to 2 when all *p*_*i*_ = 1/4 and 0 when *p*_(1)_ = 1. In blue are the entropy and corresponding values of *p*_(1)_ for all cases where MPEE selects a triply ambiguous character. In green, MPEE selects a doubly ambiguous character. In red are the cases where MPEE selects a non-ambiguous character.

The solution that is proposed here relies on augmenting the set of states the inference can choose from compared to the actual set the data generating process is based on. The objective then amounts to choosing a non-ambiguous state (i.e., a single nucleotide) whenever the signal in the data is strong, or an ambiguous one (i.e., a combination of 2, 3 or 4 nucleotides) when the signal is weak. We designed a new criterion, called the Minimum Posterior Expected Error (MPEE) that is based on the calculation of a score associated to each (ambiguous and non-ambiguous) state. This score relies on the marginal probabilities calculated for each non-ambiguous state along with error values that depend on the level of ambiguity of the ancestral state considered for the inference.

We evaluated the performance of standard criteria for the inference of ancestral sequences along with our new approach on simulated data. In particular, we focused of the traditional Maximum A Posteriori (MAP) criterion, which consists in selecting the (non-ambiguous) state with the largest marginal probability at a given node and site. The MAP criterion is that implemented in the software PAML (*Yang, 1997*). We also considered a modification of the MAP criterion, the MAP_r_ criterion, that only considers a subset of the available data to draw inference from. This criterion is implemented in RAxML. Our results indicate that MAP is less prone to errors compared to MAP_r_, although further investigations, with a wider range of simulation settings, would be needed in order to confirm the trend observed here.

MPEE outperforms the other criteria although this conclusion depends on the way errors are measured, in particular in cases where ambiguous states are selected for the inference. Also, the comparison of results obtained when the substitution model is misspecified demonstrates the robustness of this criterion, both in terms of the fraction of errors as well as the distribution of marginal probabilities of inferred states. Importantly, about half of the errors arising when the MAP criterion is used for the inference are corrected when applying the MPEE criterion instead, whether the model is correctly specified or not. This gain in terms of accuracy of the estimates comes with a modest decrease of the precision. Indeed, about 2% of the (non-ambiguous) states correctly estimated with MAP are considered as ambiguous when using MPEE.

The new criterion for ancestral state reconstruction that is presented in this study therefore helps improving the accuracy of the inference without compromising substantially on the precision of the solutions returned. It also brings forward information about the uncertainty in the inference in a concise format, which is relevant to biologists. Finally, the computational overload associated to the application of this new criterion is negligible compared to the standard approach, thereby making this new criterion a sensible alternative to the existing ones.

## Software availability

The MPEE and the MAP critera for inferring ancestral states are implemented and documented in the PhyML software package (https://github.com/stephaneguindon/phyml).

## Acknowledgements

This research was supported by the Institut Français de Bioinformatique (RENABI-IFB, Investissements d’Avenir).

